# Dravet variant *SCN1A^A1783V^* impairs interneuron firing predominantly by altered channel activation

**DOI:** 10.1101/2021.04.23.440874

**Authors:** N. Layer, L. Sonnenberg, E. Pardo González, J. Benda, H. Lerche, H. Koch, TV. Wuttke

## Abstract

Dravet syndrome (DS) is a developmental epileptic encephalopathy mainly caused by functional Na_V_1.1 haploinsufficiency in interneurons (IN). Recently, a new conditional mouse model expressing the recurrent human p.A1783V missense variant has become available. Here we provide an electrophysiological characterization of this variant in tsA201 cells, revealing both altered voltage-dependence of activation and slow inactivation without reduced sodium peak current density. Simulating IN excitability in a Hodgkin-Huxley one-compartment model suggested surprisingly similar firing deficits for *Scn1a*^A1783V^ and full haploinsufficiency as caused by heterozygous truncation variants. Impaired Na_V_^A1783V^ channel activation was predicted to have a significantly larger impact on channel function than altered slow inactivation and is therefore proposed as the main mechanism underlying IN dysfunction. The computational model was validated in cortical organotypic slice cultures derived from conditional *Scn1a^A1783V^* mice. Pan-neuronal activation of the p.A1783V variant *in vitro* confirmed the predicted IN firing deficit while demonstrating normal excitability of pyramidal neurons. Taken together these data demonstrate that despite maintained physiological peak currents density LOF gating properties may match effects of full haploinsufficiency on neuronal level, thereby causing DS.

**Highlights:**

Na_V_1.1^A1783V^ alters voltage-dependence of activation and slow inactivation while not affecting fast inactivation.
Depolarizing and hyperpolarizing shifts of activation and slow inactivation curves result in combined channel loss of function (LOF).
Simulations of Na_V_1.1^A1783V^ interneuronal properties indicate reduced action potential firing rates comparable to full *SCN1A* haploinsufficiency, which is often found in Dravet syndrome.
*In silico* modelling identifies impaired channel activation as the predominant mechanism of channel LOF.
Panneuronal induction of *Scn1a*^+/A1783V^ in a cortical slice culture model confirms restriction of loss of function and its restriction to interneurons.

## Introduction

Dravet syndrome (severe myoclonic epilepsy in infancy (SMEI)) is a rare form of developmental epileptic encephalopathy characterized by an early febrile seizure onset in childhood followed by a high frequency seizure period with large diversity between seizure types (Dravet, 1978). The epilepsy phenotype is combined with a multitude of comorbidities, including intellectual disability, sleep disorder, motor dysfunction and an increased incidence for sudden death in epilepsy patients (SUDEP). More than 80% of diagnosed Dravet cases are caused by de-novo loss of function (LOF) variants causing functional haploinsufficiency of the *SCN1A* gene coding for the alpha-subunit of the voltage-gated sodium channel Na_V_1.1 (Marini et al., 2011). Na_V_1.1 variants are associated with a variety of neurological diseases including familial febrile seizures, generalized epilepsy with febrile seizures plus (GEFS+), sporadic/familial hemiplegic migraine and Dravet syndrome resembling a clinical phenotype at the severe end of the spectrum (Gambardella & Marini, 2009). The high diversity of *SCN1A* associated disorders can be explained by a variety of gain and loss of function variants (Catterall, Kalume, & Oakley, 2010) which impair the channel on different levels including changed RNA transcription (Lange et al., 2019), reduced protein expression, altered membrane trafficking (Thompson, Porter, Kahlig, Daniels, & George, 2012), impaired β-subunit interaction (Spampanato et al., 2004) and channel dysfunction (Kluckova et al., 2020).

In the last years, different cell culture and animal models have been established based on patient mutations to explore the different pathomechanisms of Na_V_1.1 leading to epileptic phenotypes. Most frequently, heterozygous Na_V_1.1 knockout mice, mimicking protein truncating nonsense variants found in Dravet patients, are used to investigate pathophysiological disease mechanisms (Favero, Sotuyo, Lopez, Kearney, & Goldberg, 2018; Mistry et al., 2014; Ogiwara et al., 2007; Yu et al., 2006). Such studies revealed a predominant role of LOF effects in certain inhibitory neuron classes including parvalbumin interneurons (PV-IN) (Bechi et al., 2012; Favero et al., 2018; Rubinstein et al., 2015; Tai, Abe, Westenbroek, Scheuer, & Catterall, 2014; Tiraboschi et al., 2020; Tran et al., 2020) However, nonsense mutations only account for a subset of pathogenic variants leading to epileptic phenotypes: Especially milder epilepsy syndromes are often caused by missense variants within the protein coding sequence and were studied in IPSC-derived neurons (Xie et al., 2020) and mouse models (Das et al., 2021; Hedrich et al., 2014). However, also DS may be caused by missense variants (Marini et al., 2011). While protein truncation (nonsense variants) or diminished surface expression (certain missense variants) result in pronounced reduction of sodium peak currents the mechanisms of remaining missense variants can be more complex, as physiological gating properties of channels may be affected on multiple levels. While full haploinsufficiency is modelled well by heterozygous *Scn1a* knock out mice *(PMID: 23922229)*, they may not generally reflect Dravet patient disease status for carriers of missense variants. To approach this uncertainty, conditional mice expressing the human Dravet missense variant p.A1783V (Depienne et al., 2009; Klassen et al., 2014) have been recently developed by the Dravet Syndrome European Federation (PMID: 31578435). However, the effects of p.A1783V on Na_V_1.1 channel function have not been delineated to date. A limited number of studies provide first insight in the effects of variant *Scn1aA^1783V^* channels on neuron excitability after breeding conditional *Scn1aA^1783V^* mice with Cre-driver lines. Depending on the exact genetic background of offspring animals, the subtype of studied neurons and the brain areas in which neurons were recorded, differential effects on neuron firing have been described (Almog et al., 2021; Kuo, Cleary, LoTurco, Chen, & Mulkey, 2019). Although loss of function of *Scn1aA^1783V^* has been assumed, the biophysical effects of the variant on Na_V_1.1 channel function have not yet been delineated. In this study we are providing a detailed comparative characterization of variant p.A1783V and wildtype (WT) gating properties in tsA201 cells. While full haploinsufficiency with reduced Na^+^ peak currents is a common feature in DS our recordings revealed clear LOF mechanisms, albeit with preserved overall sodium peak current density. Building on these data a single compartment computational neuron model was developed to predict and compare firing deficits in inhibitory and excitatory neurons for p.A1783V in comparison to full haploinsufficiency / heterozygous knockout. The model was electrophysiologically validated in cortical organotypic brain slice cultures derived from *Scn1aA^1783V^* mice. Interestingly, despite overall normal sodium peak current density, p.A1783V was predicted to cause a fairly strong IN firing deficit comparable to the heterozygous knockout condition. As for numerous other channelopathies correlations of the nature of *SCN1A* mutations (such as location in the channel, genetic mechanism and associated impact on encoded protein function) with disease severity have been described (Zuberi et al., 2011). Our data demonstrate that altered channel gating can outweigh maintained physiological sodium peak current density and translate pronounced functional impairment on neuronal level as frequently associated with DS.

## Material and methods

### Mutagenesis

The human Nav1.1 channel cloned in pCDM8 vector as described before (Hedrich et al., 2014) was corrected for two point mutations E650V and S1969A (Peters, Rosch, Hughes, & Ruben, 2016) by site directed mutagenesis (Agilent technologies). The wildtype open reading frame included the canonical *SCN1A* adult isoform 2 (total length of 5997bp) equivalent to transcript NM_006920.5. To engineer the missense mutation p.A1783V into the human Na_V_1.1 channel, site-directed mutagenesis was performed using PCR with Pfu polymerase (Promega; mutagenesis primers 5’ to 3’; F: ATG TAC ATC GTG GTC ATC CTG GAG AAC TTC AGT, R: AGG ATG ACC ACG ATG TAC ATG TTC ACC ACA). The introduced mutations were verified and further mutations were excluded by sequencing the whole *SCN1A* cDNA prior to using the clones for transfection. Plasmid purification was performed from E.Coli One Shot TOP10/P3 (Thermo Scientific).

### tsA201 cell culture and transfection

tsA201 cells were cultured in Dulbecco’s modified Eagle nutrient medium (Gibco) supplemented with pyruvate, 10% v/v fetal bovine serum (Pan Biotech) and 2 mM L-glutamine (Biochrom) at 37°C in a 5% CO_2_ humidified atmosphere. For transfection, cells with a passage from P15 - P22 were used. Before transfection, 800.000 - 1.000.000 cells were split in 35 mm petri dishes. 6 hours later, cells were transfected following standard transfection protocols for Mirus Trans-IT transfection agent: 4 μg of wildtype or mutant human *SCN1A* cDNA, encoding the Na_V_1.1 channel-α-subunit, and each of 0.4 μg of the human β_1_- and β_2_-subunits of voltage-gated Na_V_ channels, which had been previously modified to express either green fluorescent protein (pCLH-hb1-EGFP) or a CD8 marker (pCLH-hb2-CD8) to label cells expressing both subunits (Liao et al., 2010), were added to 250 μl of Opti-MEM (Gibco) and 7.5 μl of Mirus transit agent (Mirus Bio). After 20 minutes, transfection mixture was added to the cells. Electrophysiological whole cell recordings were performed 48 hours after transfection from tsA201 cells expressing all three sodium channel subunits indicated by an inward sodium current (α-subunit), surface coting with anti-CD8 antibody coated microbeads (Dynabeads M450, Dynal) (β_1_-subunit) and green fluorescence (β_2_-subunit).

### Cortical brain slice cultures

Coronal brain slices from postnatal *B6(Cg)-Scn1a^tm1.1Dsf^*/J mice (Jackson laboratories) of either sex were obtained at P4 - 5. Mice were quickly decapitated and the cranium, brain stem and hindbrain were removed. The forebrain was placed in ice-cold artificial cerebrospinal fluid (aCSF) bubbled with carbogen (95% O_2_/5% CO_2_). The aCSF contained [in mM]: 118 NaCl, 3 KCl, 1.5 CaCl_2_, 1 MgCl_2_, 25 NaHCO_3_, 1 NaH_2_PO_4_, 30 glucose, pH 7.4 with an osmolarity of 310-320 mOsm/kg. Forebrains were glued with cerebellar end facing downwards onto an agar block and 350 μm-thick slices were cut using a vibratome (Microm, H650 V) in ice-cold aCSF. Afterwards, slices were placed in 36° C warm aCSF (bubbled with carbogen) for 10 minutes. Subsequently, coronal slices containing the somatosensory cortex were transferred to Millicell cell culture inserts (Merck Millipore) floating on slice culture medium (Minimum Essential Medium Eagle with 20% horse serum, 1 mM L-Glutamine, 0.00125% ascorbic acid, 0,001 mg/ml insulin, 1mM CaCl_2_, 2 mM MgSO_4_, 1% Penicillin/Streptomycin,13 mM Glucose, pH 7,28 and osmolarity 320 mOsm/kg).

### Electrophysiological recordings in tsA cells

Whole cell patch clamp recordings in tsA201 cells were performed 48 hours after transfection. Cells were split in 35 mm petri dishes 1 hour prior to recordings and cells in each dish were used for up to one hour after transfer to the patch clamp setup. The extracellular bath solution contained [in mM]: 140 NaCl, 4 KCl, 1 MgCl_2_, 2 CaCl_2_, 5 Hepes and 4 Glucose. pH was adjusted with HCl to 7.4 and osmolarity was 300-305 mOsm/kg. Patch pipettes were pulled from borosilicate glass (Science Products GmbH) using a Sutter P97 Puller (Sutter Instruments), with resistances of 1.5 – 3.0 MΩ. Intracellular solutions for patch pipettes contained [in mM]: 5 NaCl, 2 MgCl_2_, 5 EGTA, 10 Hepes and 130 CsF, pH of 7.4 (adjusted with CsOH) and an osmolarity of 290-295 mOsm/kg. Before seal formation cells were incubated for one minute with with anti-CD8 antibody coated microbeads. Signals were amplified with an Axopatch 200B (Molecular Devices) amplifier, digitized by a DigiData 1440 digitizer (Molecular Devices) and recorded with pClamp 10.7 software (Molecular devices). During recordings cells were held at −120 mV.

### Electrophysiological recordings in slice culture

Whole cell patch clamp recordings of excitatory pyramidal cells and fast-spiking inhibitory neurons in the somatosensory cortex were acquired 7-14 days after viral transduction with AAV8-hSyn-Cre-GFP (SignaGen) in organotypic cortical slice culture using an Axopatch 200B (Molecular Devices) amplifier, a DigiData 1440 digitizer and pClamp 10.7 software. Slices were positioned in a submerged-type recording chamber (Luigs & Neumann), continuously superfused with oxygenated recording aCSF and maintained at a temperature of 33 ± 1 °C. Cortical neurons transduced with AAV8-hSyn-Cre-GFP were visualized with an Axioskop 2FS (Zeiss) microscope. Patch pipettes were pulled from borosilicate glass (using a Sutter P97 Puller) with resistances of 2.5 – 5.5 MΩ. Intracellular solutions for patch pipettes contained [in mM]: 140 K-Gluconate, 1 CaCl_2_, 10 EGTA, 2 MgCl_2_, 4 Na_2_-ATP, 10 HEPES and 0.45% biocytin, pH of 7.2 and an osmolarity of 300-310 mOsm/kg. Cortical pyramidal cells and fast spiking cortical inhibitory neurons were used for current clamp recordings. Cell identity was determined via morphological and electrophysiological properties and by *posthoc* GAD67-staining after slice fixation. Whole-cell recordings were compensated for cell capacitance and series resistance 5 min after rupturing the seal. Recordings were corrected for a liquid junction potential of 15 mV and held at −70 mV. To block postsynaptic AMPA receptor driven depolarization waves, 20 μM CNQX (Sigma Aldrich) was added to the bath solution prior to recording. Cells showing unstable series resistance, resting membrane potential or shifts in resting membrane potentials were excluded from analysis. The input resistance of transfected cortical neurons was determined by the slope of a linear regression fit to corresponding steady state voltage responses plotted versus a series of current injections ranging from −10 to −110 pA with −10 pA increments. For recording of action potentials, only events with a voltage peak amplitude surpassing 0 mV were regarded as action potentials. Neuronal train firing properties were analyzed by current squared pulse injections of increasing intensity (starting at – 50 pA and increasing by 25 pA per sweep up to +300 pA current injection for pyramidal cells; starting at – 0 pA and increasing by 50 pA per sweep up to +700 pA current injection for fast spiking interneurons) for 800 ms followed by a 5 s inter-sweep interval before the following current injection.

### *In silico* modelling of cortical neurons

Simulations were run with custom Python 3.7 software. We used a single compartment-based model with a cylindrical shape with length L and diameter d. The model is based (Pospischil et al., 2008) and consists of two sodium currents *I_Na,wt_* and *I_Na,mut_*, a delayed rectifier potassium current *I_K_*, a M-type potassium current *I_M_* and a leak current *I_leak_* with

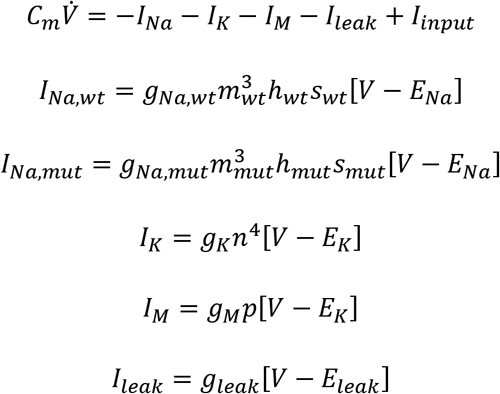

with membrane capacitance *C_m_* = 1μ*F*/cm^2^. *I_input_* the input current, *I_i_* are the ionic currents with maximal conductance *g_i_*, reversal potential *E_i_* and gating variables *m, h, n, p* with dynamics

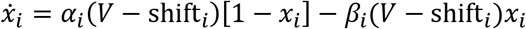

and the slow inactivating gating variable *s* with

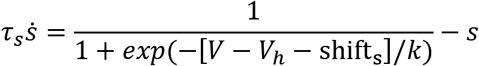

where *ẋ_i_* is the derivative of the gating parameter with respect to time, α_*i*_(*V*) is the opening rate and β_*i*_(V) the closing rate of the respective gate *x_i_*. Steady-state curves and time constants are given by *x*_∞,*i*_(V) = α_*i*_(*V*)/[α_*i*_(*V*) + β_*i*_(*V*)] and τ_*i*_(V) = 1/[α_*i*_(*V*) + β_*i*_(*V*)], respectively. The dynamics of the slow inactivating gating variable *s* followed

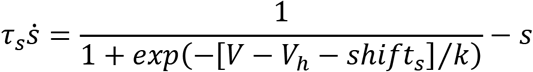

Like in Pospischil et al. (2008), parameters for gating variables *m*, *h* and *n* are taken from Traub and Miles (1991) and for gating variable p from Yamada (1989). We constructed the slow inactivation gate s to be similar to observed kinetics in Vilin and Ruben, (2001). To simulate different neuron types, we adapted the conductivities of the ionic currents Pospischil et al. (2008). All parameters that differ from Pospischil et al. (2008) are summarized in **Table 1**. Effects of mutations were simulated with changes in the shifting parameter *shift_i_* for the activation and slow inactivation sodium gates with *shift_i_* = 0 unless stated otherwise.

**Table 1:**
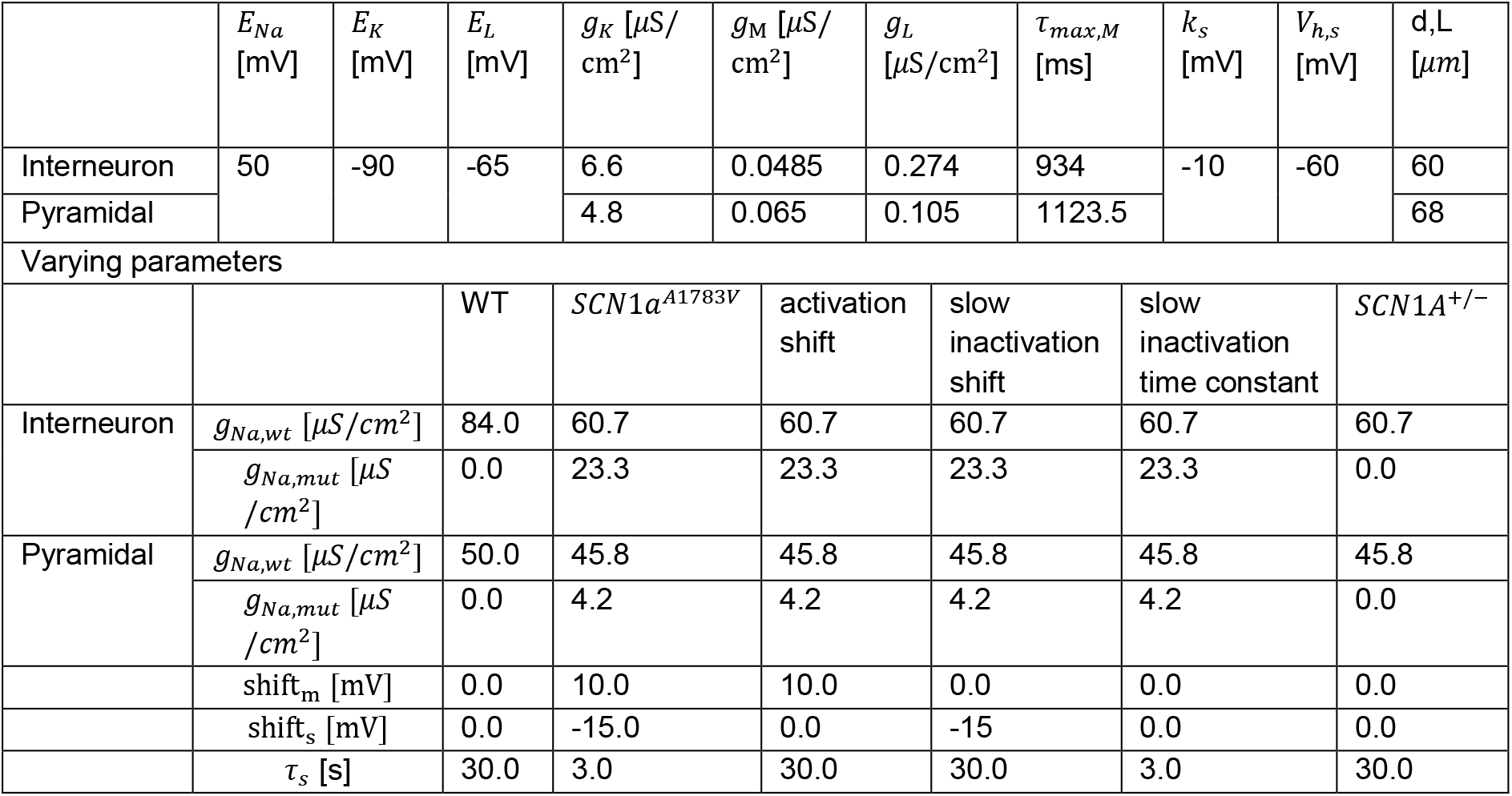
Simulation parameters for the cortical neuron models.

To simulate the effects of heterozygous mutations in *SCN1A* on the firing behaviour of pyramidal neurons and interneurons, we modelled different ratios of *SCN1A* and *SCN8A* expression in the respective neuron types with the assumption of equal gating kinetics in both sodium channels (*g_Na,wt_* = *g_SCN1A_*/2 + *g_SCN8A_* and *g_Na,mut_* = *g_SCN1A_*/2). The ratio of gene expression was *g_SCN8A_* = 5*g_SCN1A_* for pyramidal neurons and *g_SCN8A_* = 0.8*g_SCN1A_* for interneurons based on expression data from cortical mouse neurons (Yao et al., 2020). To investigate how each observed change in gating parameters contributes to the observed change in firing behaviour, we constructed several mutation models, one combined mutation model with all parameter changes and for each adapted parameter one model with only one respective change.

### Whole cell patch clamp data analysis

Currents for activation and inactivation properties of Nav1.1 sodium channel in expressed in tsA201 cells were recorded as described before (Hedrich et al., 2014). Conductance was determined by plotting the observed peak Na^+^-current and fitting by

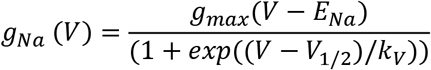

with *I* as peak current, *V* as command voltage, *g_Na_* being the conductance, *g_max_* the maximal conductance, *E_Na_* the observed reversal potential, *k_V_* as slope factor and *V_1/2_* the half maximal activation. Steady-state fast and slow inactivation were fit to the Boltzmann equation:

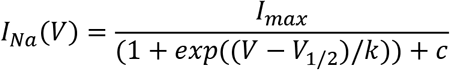

with *I_max_* as the maximal evoked sodium current and *c* being an additional constant. Recovery from fast inactivation was analyzed by fitting with a 1-exponential equation:

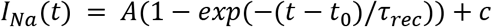

with *A* as the maximal recovered current, t_0_ as the initial time delay and *τ_rec_* as the time constant of recovery from fast inactivation.

Entry into slow inactivation was fit by

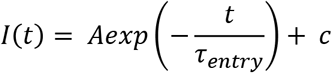

with *I*(*t*) as remaining peak current after various duration of depolarizing pre-pulse, *A* as maximal amplitude of peak current, *t* the duration of pre-pulse and *τ_entry_* as the time constant of slow inactivation.

Use dependence was fit by a second-order exponential equation with

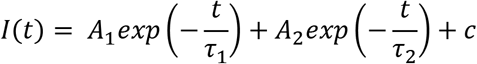

Single action potentials properties for current clamp whole-cell recordings in slice culture were analyzed as followed: Input resistance was determined by the linear regression of plateau potentials at the end of a 500 ms long square pulse with negative current injection from −10 to −120 pA. Active properties of action potentials were analyzed from the second action potential in train at a step current injection of 600 pA for fast-spiking interneurons and at 100 pA for pyramidal cells. The threshold of an action potential was determined as the voltage at which the first derivative *dV/dt* reached 20 mV/s. Action potential amplitude and rise time were measured from the threshold to the peak and the half-width was determined as the duration from 50% of the peak amplitude from the rising to the corresponding potential of the falling phase.

### Statistical analysis

All data are displayed as Mean ± SEM. Statistical testing was performed via unpaired, two-tailed t-test, *n* indicates the number of cells.

All data were analyzed using Clampfit software of pClamp 10.703 (Axon Instruments), Microsoft Excel (Microsoft Corporation, Redmond, WA, USA). Statistics were performed using Graphpad 7 software (Graphpad prism, San Diego, CA, USA).

## Results

### Biophysical characterization of *SCN1A^A1783V^* channel function in tsA201 cells reveals LOF with maintained peak sodium currents

Variant or WT channels were transfected into tsA201 cells and whole-cell patch clamping was performed Representative raw current traces are shown in **Figure 1A**. While peak current density was not changed for variant channels in comparison to WT (**Figure 1B**) activation and inactivation properties were markedly altered. We found a significant right shift of the half-maximal conductance indicating that variant channels open and reach their maximal activation at more depolarized potentials (**Figure 1C**). Fast inactivation was unaffected as revealed by comparable voltage-dependence of steady state fast inactivation (**Figure 1B**), time constant *τ* (**Figure 1E**) and recovery from fast inactivation (**Figure 1D**) for WT and Na_V_1.1^A1783V^ channels. In contrast, entry into slow inactivation was accelerated (**Figure 1G**) and voltage-dependence was shifted to hyperpolarized potentials (**Figure 1H**). Overall loss of Na_V_1.1^A1783V^ channel function became very obvious upon rapid successive depolarizing stimulations at 40 Hz revealing a markedly increased use-dependence with consecutively pronounced run down of the sodium current (**Figure 1F**).

**Figure 1:**
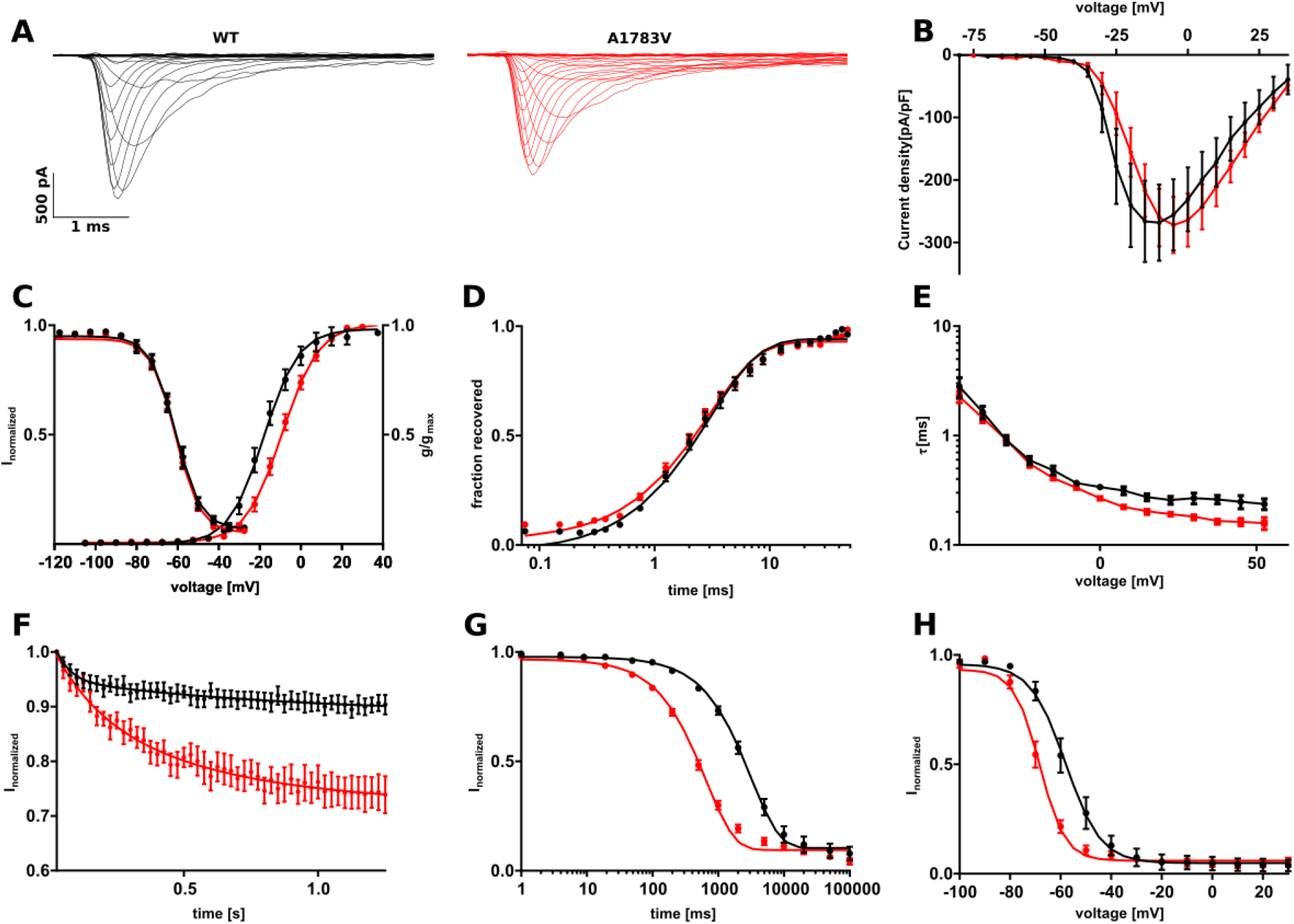
Biophysical properties of *SCN1A* wildtype and mutant channels *SCN1A^A1783V^* recorded in tsA201 cells display a clear loss of function. (A) Representative currents recorded from tsA201 cells transfected with *SCN1A^WT^* (black traces) or *SCN1A^A1783V^* (red traces). (B) Peak Na^+^ currents normalized by cell capacitance were plotted versus command voltage and revealed no changes between WT and variant *SCN1A^A1783V^* channels. (C, right curve) Voltage-dependant steady state activation curves represented as normalized conductance fit by a Boltzmann function to the data points. (C, left curve) Voltage-dependent state of fast inactivation fit by a Boltzmann function to the data points. (D) Time course of recovery from fast inactivation at −100 mV. (E) Voltage-dependence of the time constant of fast inactivation *τ_h_* analysed by a one exponential fit to inactivating phase of the sodium currents from the activation protocol. (G) Entry into slow inactivation represented as a first-order exponential fit to the data points. (H) Voltage-dependent steady-state of slow inactivation fit by a Boltzmann function as in (C). (F) Sodium current use dependence evoked by 50 pulses at a frequency of 40 Hz as normalized peak currents plotted against the time fit by a second order exponential fit. Number of recorded cells, exact values of statistically analysed parameters and p-values are listed in **Supplement table 1**.

### *In silico* modelling reveals pronounced interneuron firing impairment by Na_V_1.1^A1783V^

Full haploinsufficiency is considered the genetic mechanism underlying the majority of *SCN1A* Dravet variants. About half of the described variants lead to protein truncation with reduced protein expression in affected neurons and likely markedly reduced current conductance (Marini et al., 2011). Our recordings revealed clear biophysical LOF changes of Na_V_1.1^A1783V^ compared to WT, however no reduction of peak current density. Therefore, we next asked how these combined functional alterations translate to neuronal level and how they compare to the heterozygous knock out condition. To address these questions we built a Hodgkin-Huxley single compartment conductance-based model with the combined kinetic alterations reflecting intrinsic and firing properties of either cortical fast-spiking interneurons (IN) or excitatory neurons (EN). Differential *SCN1A* to *SCN8A* expression ratios of IN and EN as well as either the WT (Na_V_1.1^+/+^), the variant (Na_V_1.1^+/A1783V^) or the full haploinsufficiency (Na_V_1.1^+/-^) condition were implemented. The parameters for the different model types are summarized in **Table 1**. Additionally, we simulated altered voltage-dependence of activation as well as altered voltage-dependence and kinetics of slow inactivation separately to dissect their joint or exclusive impact on neuron action potential firing in comparison to the WT and the full haplonisufficiency condition **(Figure 2A)**.

**Figure 2:**
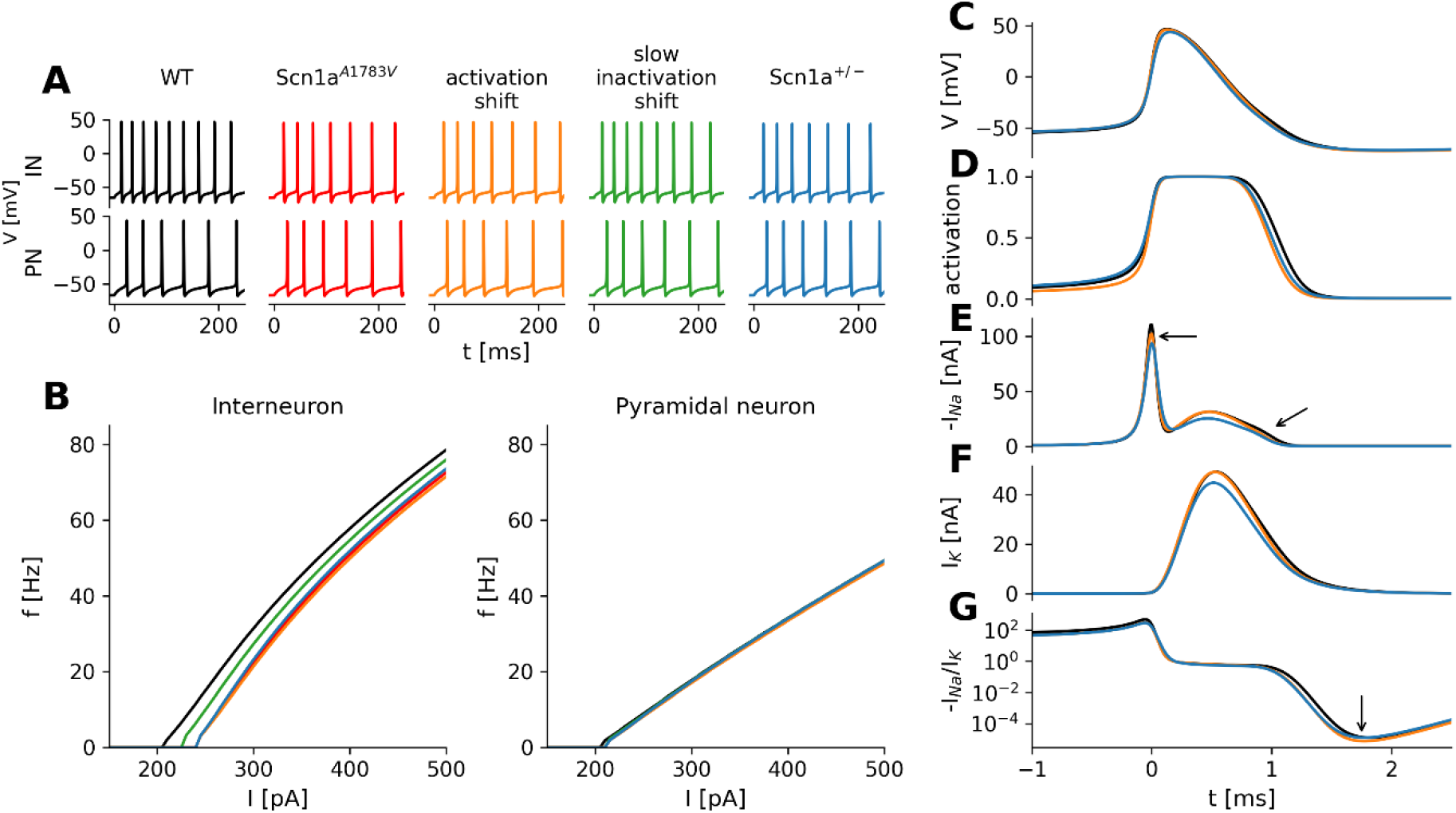
*In silico* modelling of neuron excitability. (A) Action potentials in response to square pulse current injections of 300 pA of simulated cortical fast–spiking interneurons (upper traces) and cortical pyramidal cells (lower traces) with various modifications of Na_V_1.1 current kinetics based on findings in tsA recordings. Colors indicate different model conditions: WT (black), heterozygous shift of the voltage-dependence of steady-state activation (orange), heterozygous shift of the voltage-dependence of the steady-state slow inactivation (green), combination of both changes (red) and a heterozygous protein knockout (blue). The exact parameters for all simulated conditions are summarized in **Table 1**. (B) FI-curves of interneurons (left) and pyramidal cells (right) constructed from simulation studies displayed in (A). (C-G) Time courses of model variables during the first action potential after a 300 pA current injection. (C) Membrane voltage; (D) ratio of open sodium channels; (E) negative of sodium current; (F) potassium current; (G) absolute ratio of sodium current to potassium current.

In the interneuron model, solely shifting the activation curve to more depolarized membrane potentials caused the most severe reductions of the firing rates in comparison to the WT in the firing behaviour, closely followed by the *SCN1A^A1783V^* and *SCN1A^+/^* conditions. Shifting the slow inactivation curve to more hyperpolarized potentials reduced firing rates to a lesser extend **(Figure 2B, left)**, whereas the faster slow inactivation model showed almost no changes **(Supplement figure 1A,B)**. Simulations of the model with faster slow inactivation showed almost no difference to the WT firing rates, but were the only simulation with a similar accelerated rundown of sodium current amplitude as observed in TsA201 cells **(Supplemental Figure 1C)**. All of these changes had almost no effect on the firing rate of the pyramidal model **(Figure 2B, right)**.

To better understand why shifting the activation curve for 50% of the Na_V_1.1. channels (to mimic the heterozygote patient situation) had a stronger effect on neuronal firing than the simulation of the heterozygous protein knock-out, we analyzed the dynamics of the first action potential waveform in more detail (**Figure 2C-G)**. Although on a first glance the action potential waveforms seemed to be rather similar (**Figure 2C**), the sodium activation gate (**Figure 2D**) of the model with a shifted activation curve (orange) opens later and closes earlier than of the WT (black) and the *SCN1A^+/-^* models (blue). The resulting sodium current is reduced during AP initiation and rising phase of the action potential compared to the WT simulation but is similar during the falling phase of the action potential in comparison to the WT simulation. Only at the very end of the AP is the sodium current is again slightly reduced. The *SCN1A^+/-^* model in contrast has a generally reduced sodium current (**Figure 2E**). The delayed rectifier potassium current is affected in similar ways by the variants. For the *SCN1A^+/-^* model its reduction is most prominent (**Figure 2F**). However, relative to the potassium current the sodium current is most strongly reduced by the shift in the activation curve found during the final phase of afterhyperpolarization (**Figure 2G**).

### *Scn1a^A1783V^ LOF* is restricted to *inhibitory neurons in cortical mouse brain slice cultures*

To confirm the simulated effects of *SCN1A^A1783V^* on a neuronal firing, we performed whole-cell patch clamp recordings in cortical brain slice cultures derived from P4-5 heterozygous B6(Cg)-*Scn1a^tm1.1Dsf^*/J mice or WT littermates for control. Cultures were transduced at day 1 *in vitro* (1DIV) with adeno-associated-virus 8 (AAV8) encoding Cre-recombinase under the human synapsin promoter in order to induce recombination and subsequent expression of the p.A1783V variant. Since *Scn1a* is upregulated in the postnatal period only from P11 onwards (Cheah et al., 2013), this early expression of Cre recombinase allowed for recombination of transduced neurons carrying the floxed *Scn1a^A1783V^* allele before endogenous upregulation of the *Scn1a* gene. This approach ensured activation of the variant following the endogenous expression time-course of the Na_V_1.1 channel. 7 to 14 days after successful viral transduction indicated by the fluorescent reporter EGFP, we patched transfected cortical neurons: We found a loss of function in cortical fast-spiking interneurons (**Figure 3D**) while the firing of cortical excitatory neurons (**Figure 3B**) was not altered. Transduced fast-spiking interneurons of floxed mice displayed a significantly increased rheobase, a reduced maximum firing frequency as well as a reduced input resistance compared to transduced interneurons of WT littermates. For other active single AP properties, we didn’t find any alterations; AP threshold, AP amplitude, AP rise time and AP half width within evoked AP trains were undistinguishable between transduced WT and heterozygous mutant slices. This LOF of neuronal firing found in fast-spiking interneurons was not present in patched cortical pyramidal neurons as indicated by unaltered f-I curves of AP trains. Nevertheless we could detect a small reduction of the input resistance of EN. However, this effect was too mild to cause alterations of the analysed action potential properties.

**Figure 3:**
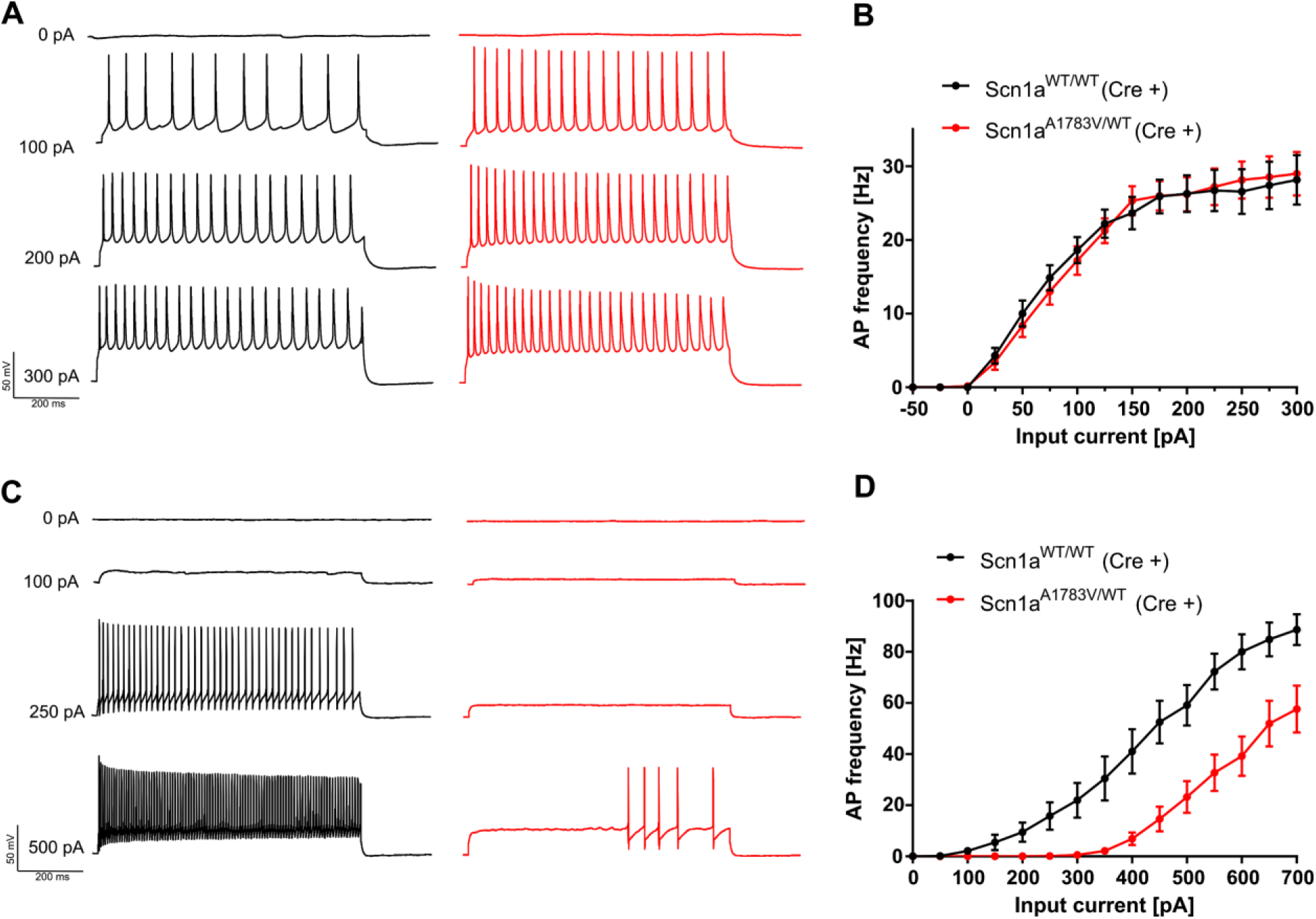
In vitro studies of neuronal excitability. (A) Representative AP trains in response to 800 ms step current injections from cortical pyramidal cells in brain slice cultures and (**C**) from fast-spiking interneurons expressing WT (black traces) and heterozygous variant Na_V_1.1^+/A1783V^ channels (red traces). (**B**) f-I curves of pyramidal cells from heterozygously floxed mice showed the same frequency of action potential firing while (**D**) fast-spiking inhibitory neurons started eliciting action potentials at higher current injections and maintained lower AP frequencies up to higher current injections. Number of recorded cells, exact values of statistically analyzed parameters and p-values are listed in **Supplemental Tables 2 & 3**.

## Discussion

Full haploinsufficiency is considered the leading genetic cause of DS, resulting in impaired excitability predominantly of interneurons (Catterall, Dib-Hajj, Meisler, & Pietrobon, 2008). In this study we functionally characterized the human recurrent *Scn1a^A1783V^* DS variant. While reduced sodium peak current density is a common feature associated with DS, our analysis revealed altered activation and slow inactivation properties, albeit with preserved sodium peak current density. However, an enhanced use-dependent rundown of Na^+^ current was observed upon repetitive stimulations. Although an interference of the variant (localized at the end of segment 6 in domain 4) with local determinants of fast inactivation would have been conceivable (Ulbricht, 2005) fast inactivation was found unchanged. Since fast inactivation was unaffected, use-dependence can only be attributed to slow inactivation characteristics of the variant. This could be indeed be confirmed by *in silico* modelling of Na^+^ current with physiological or accelerated slow inactivation kinetics **(Supplemental Figure 1C).** Interestingly modelling of neuron excitability predicted a fairly strong overall interneuron AP firing deficit for Na_V_1.1^+/A1783V^ at comparable levels to the simulated full haploinsufficiency condition (Na_V_1.1^+/-^). Since our tsA201 cell recordings had revealed a ~25 % sodium current rundown upon 40 Hz stimulation we speculated that altered slow inactivation and associated enhanced use dependence might largely account for the predicted interneuron firing deficit. Importantly the model correctly predicted enhanced use-dependence of Na_V_1.1^*A1783V*^ currents and confirmed accelerated slow inactivation as the only underlying mechanism **(Supplemental Figure 1C)**. However, when simulating f-I curves for fast spiking interneurons with different time constants of slow inactivation, an acceleration of *τ* only had a marginal effect on neuronal firing **(Supplement Figure 1A)**. Interestingly *in silico* modelling predicted a fairly strong overall interneuron AP firing deficit for Na_V_1.1^+/A1783V^ at comparable levels to the full haploinsufficiency condition (Na_V_1.1^+/-^).

According to our simulations, shifted voltage-dependence of fast activation was predicted as the main driver of the impaired interneuron firing. Comparison to the simulated Na_V_1.1^+/-^ condition revealed a surprisingly similar action potential deficit. While surprising at first these findings could at least partially be explained when putting the activation properties and relative levels of conducted currents of Na_V_1.1^+/*A1783V*^ and Na_V_1.1^+/-^ in context with the different phases of the action potential. Conductance based modelling suggested a reduced number of sodium channels available to participate in action potential initiation due to the shifted activation curve and subsequently delayed opening of the channels. This effect was qualitatively similar to Na_V_1.1^+/-^ (with only a limited number of available channels due to the simulated heterozygous knock out condition) in this early phase of an action potential and was predicted to account for most of the overall LOF. In the falling phase of the action potential, however, Na_V_1.1^+/*A1783V*^ yielded relatively larger currents than Na_V_1.1^+/-^ and accordingly recruited more potassium channels. Our simulations predicted that the induced potassium current outweighed the conducted sodium current in the early hyperpolarzing phase in neurons expressing Na_V_1.1^+/A1783V^. This relative surplus of K^+^ current led to pronounced afterhyperpolarization and a prolonged time window until the next spike could occur, altogether increasing the LOF effect. Consistent with our data pronounced shifts of voltage-dependence of activation have also been described for other LOF mutations associated with epilepsy, such as *SCN1A^I1656M^* (Lossin et al., 2003), *SCN1A^D249E^* (Kluckova et al., 2020) and *SCN1A^R859C^* (Barela et al., 2006). Like *SCN1A^A1783V^* these variants also feature unaltered current density in comparison to the WT. In future studies it would be interesting to model neuron firing properties of these variants for comparison with our *Scn1a^+/A1783V^* and *Scn1a^+/-^* data and for correlation with associated clinical phenotypes.

*Scn1a* is predominantely expressed in IN (Yao et al., 2020) while EN mainly rely on Na_V_1.6 for action potential initiation. Hence a dysfunction of IN but not EN was expected and was indeed reflected *in silico*. These data were confirmed in cortical brain slice cultures derived from conditional *Scn1a^+/A1783V^* mice after Cre-dependent activation of the variant (via AAV mediated transduction of neurons within the slice cultures). We found a significant rightward shift of the FI-curve and prominent shift of the rheobase in cortical IN, while no alterations could be detected in EN. Our findings are consistent with data obtained in hippocampal interneurons expressing *Scn1a^+/A1783V^* mice which also displayed a shift in rheobase and a firing deficit at higher current injections (Almog et al., 2021).

In summary, *SCN1A^A1783V^* results in altered voltage-dependence of activation and slow inactivation while maintaining sodium peak current density. IN excitability was simulated *in silico* by a Hodgkin-Huxley one-compartment model suggesting largely similar firing impairment for Na_V_1.1^+/A1783V^ in comparison to Na_V_1.1^+/-^ resembling full haploinsufficiency. Impaired Na_V_^A1783V^ voltage-dependent activation was suggested as main mechanism underlying IN dysfunction.

## Author contribution

NL did molecular biology, cultured brain slices and performed current-clamp in brain slice culture, NL and EPG performed voltage-clamp recordings in tsA201 cells, LS and JB developed *in silico* model, NL and LS did data analysis, NL, LS, JB, HK, TVW wrote the manuscript. Proof reading and study design: all authors

## Acknowledgement

This study was supported by the German Research Foundation (DFG/FNR INTER research unit FOR2715 grants Ko4877/3-1, Le1030/15-1, He8155/1-1). TW was supported by an intramural Clinician Scientist Fellowship granted by the Faculty of Medicine, University of Tübingen (419-0-0).

## References

Almog, Y., Fadila, S., Brusel, M., Mavashov, A., Anderson, K., & Rubinstein, M. (2021). Developmental alterations in firing properties of hippocampal CA1 inhibitory and excitatory neurons in a mouse model of Dravet syndrome. Neurobiology of Disease, 148, 105209. https://doi.org/10.1016/j.nbd.2020.105209

Barela, A. J., Waddy, S. P., Lickfett, J. G., Hunter, J., Anido, A., Helmers, S. L., … Escayg, A. (2006). An epilepsy mutation in the sodium channel SCN1A that decreases channel excitability. Journal of Neuroscience, 26(10), 2714–2723. https://doi.org/10.1523/JNEUROSCI.2977-05.2006

Bechi, G., Scalmani, P., Schiavon, E., Rusconi, R., Franceschetti, S., & Mantegazza, M. (2012). Pure haploinsufficiency for Dravet syndrome Na V1.1 (SCN1A) sodium channel truncating mutations. Epilepsia, 53(1), 87–100. https://doi.org/10.1111/j.1528-1167.2011.03346.x

Catterall, W. A., Dib-Hajj, S., Meisler, M. H., & Pietrobon, D. (2008). Inherited neuronal ion channelopathies: New windows on complex neurological diseases. Journal of Neuroscience, 28(46), 11768–11777. https://doi.org/10.1523/JNEUROSCI.3901-08.2008

Catterall, W. A., Kalume, F., & Oakley, J. C. (2010). NaV1.1 channels and epilepsy. Journal of Physiology, 588(11), 1849–1859. https://doi.org/10.1113/jphysiol.2010.187484

Das, A., Zhu, B., Xie, Y., Zeng, L., Pham, A. T., Neumann, J. C., … O’dowd, D. K. (2021). Interneuron dysfunction in a new mouse model of scn1a gefs+. ENeuro, 8(2), 1–16. https://doi.org/10.1523/ENEURO.0394-20.2021

Depienne, C., Trouillard, O., Saint-Martin, C., Gourfinkel-An, I., Bouteiller, D., Carpentier, W., … LeGuern, E. (2009). Spectrum of SCN1A gene mutations associated with Dravet syndrome: analysis of 333 patients. J Med Genet, 46(3), 183–191. https://doi.org/10.1136/jmg.2008.062323

Dravet, C. (1978). [Les epilepsies graves de l’enfant]. Vie Medicale, 8, 543–548.

Favero, M., Sotuyo, N. P., Lopez, E., Kearney, J. A., & Goldberg, E. M. (2018). A Transient Developmental Window of Fast-Spiking Interneuron Dysfunction in a Mouse Model of Dravet Syndrome. The Journal of Neuroscience, 38(36), 7912–7927. https://doi.org/10.1523/jneurosci.0193-18.2018

Gambardella, A., & Marini, C. (2009). Clinical spectrum of SCN1A mutations. Epilepsia, 50(SUPPL. 5), 20–23. https://doi.org/10.1111/j.1528-1167.2009.02115.x

Hedrich, U. B., Liautard, C., Kirschenbaum, D., Pofahl, M., Lavigne, J., Liu, Y., … Lerche, H. (2014). Impaired action potential initiation in GABAergic interneurons causes hyperexcitable networks in an epileptic mouse model carrying a human Na(V)1.1 mutation. J Neurosci, 34(45), 14874–14889. https://doi.org/10.1523/JNEUROSCI.0721-14.2014

Klassen, T. L., Bomben, V. C., Patel, A., Drabek, J., Chen, T. T., Gu, W., … Goldman, A. M. (2014). High-resolution molecular genomic autopsy reveals complex sudden unexpected death in epilepsy risk profile. Epilepsia, 55(2), 6–12. https://doi.org/10.1111/epi.12489

Kluckova, D., Kolnikova, M., Lacinova, L., Jurkovicova-tarabova, B., Foltan, T., Demko, V., … Soltysova, A. (2020). OPEN A Study among the Genotype, Functional Alternations, and Phenotype of 9 SCN1A Mutations in Epilepsy Patients. Scientific Reports, 1–13. https://doi.org/10.1038/s41598-020-67215-y

Kuo, F.-S., Cleary, C. M., LoTurco, J. J., Chen, X., & Mulkey, D. K. (2019). Disordered breathing in a mouse model of Dravet syndrome. In eLife (Vol. 8). https://doi.org/10.7554/elife.43387

Lange, I. M., Weuring, W., ‘t Slot, R., Gunning, B., Sonsma, A. C. M., McCormack, M., … Koeleman, B. P. C. (2019). Influence of common *SCN1A* promoter variants on the severity of *SCN1A* -related phenotypes. Molecular Genetics & Genomic Medicine, (March), e727. https://doi.org/10.1002/mgg3.727

Liao, Y., Anttonen, A. K., Liukkonen, E., Gaily, E., Maljevic, S., Schubert, S., … Lehesjoki, A. E. (2010). SCN2A mutation associated with neonatal epilepsy, late-onset episodic ataxia, myoclonus, and pain. Neurology, 75(16), 1454–1458. https://doi.org/10.1212/WNL.0b013e3181f8812e

Lossin, C., Rhodes, T. H., Desai, R. R., Vanoye, C. G., Wang, D., Carniciu, S., … George, A. L. (2003). Epilepsy-Associated Dysfunction in the Voltage-Gated Neuronal Sodium Channel SCN1A. Journal of Neuroscience, 23(36), 11289–11295. https://doi.org/10.1523/jneurosci.23-36-11289.2003

Marini, C., Scheffer, I. E., Nabbout, R., Suls, A., De Jonghe, P., Zara, F., & Guerrini, R. (2011). The genetics of Dravet syndrome. Epilepsia, 52(SUPPL. 2), 24–29. https://doi.org/10.1111/j.1528-1167.2011.02997.x

Mistry, A. M., Thompson, C. H., Miller, A. R., Vanoye, C. G., George, A. L., & Kearney, J. A. (2014). Strain- and age-dependent hippocampal neuron sodium currents correlate with epilepsy severity in Dravet syndrome mice. Neurobiology of Disease, 65, 1–11. https://doi.org/10.1016/j.nbd.2014.01.006

Ogiwara, I., Miyamoto, H., Morita, N., Atapour, N., Mazaki, E., Inoue, I., … Yamakawa, K. (2007). Nav1.1 localizes to axons of parvalbumin-positive inhibitory interneurons: a circuit basis for epileptic seizures in mice carrying an Scn1a gene mutation. J Neurosci, 27(22), 5903–5914. https://doi.org/10.1523/JNEUROSCI.5270-06.2007

Peters, C., Rosch, R. E., Hughes, E., & Ruben, P. C. (2016). Temperature-dependent changes in neuronal dynamics in a patient with an SCN1A mutation and hyperthermia induced seizures. Scientific Reports, 6(September), 1–12. https://doi.org/10.1038/srep31879

Pospischil, M., Toledo-Rodriguez, M., Monier, C., Piwkowska, Z., Bal, T., Frégnac, Y., … Destexhe, A. (2008). Minimal Hodgkin-Huxley type models for different classes of cortical and thalamic neurons. Biological Cybernetics, 99(4–5), 427–441. https://doi.org/10.1007/s00422-008-0263-8

Rubinstein, M., Westenbroek, R. E., Yu, F. H., Jones, C. J., Scheuer, T., & Catterall, W. A. (2015). Genetic background modulates impaired excitability of inhibitory neurons in a mouse model of Dravet syndrome. Neurobiology of Disease, 73, 106–117. https://doi.org/10.1016/j.nbd.2014.09.017

Spampanato, J., Kearney, J. A., De Haan, G., McEwen, D. P., Escayg, A., Aradi, I., … Meisler, M. H. (2004). A novel epilepsy mutation in the sodium channel SCN1A identifies a cytoplasmic domain for β subunit interaction. Journal of Neuroscience, 24(44), 10022–10034. https://doi.org/10.1523/JNEUROSCI.2034-04.2004

Tai, C., Abe, Y., Westenbroek, R. E., Scheuer, T., & Catterall, W. A. (2014). Impaired excitability of somatostatin- and parvalbumin-expressing cortical interneurons in a mouse model of Dravet syndrome. Proceedings of the National Academy of Sciences of the United States of America, 111(30), 3139–3148. https://doi.org/10.1073/pnas.1411131111

Thompson, C. H., Porter, J. C., Kahlig, K. M., Daniels, M. A., & George, A. L. (2012). Nontruncating SCN1A mutations associated with severe myoclonic epilepsy of infancy impair cell surface expression. Journal of Biological Chemistry, 287(50), 42001–42008. https://doi.org/10.1074/jbc.M112.421883

Tiraboschi, E., Martina, S., van der Ent, W., Grzyb, K., Gawel, K., Cordero-Maldonado, M. L., … Esguerra, C. V. (2020). New insights into the early mechanisms of epileptogenesis in a zebrafish model of Dravet syndrome. Epilepsia, 61(3), 549–560. https://doi.org/10.1111/epi.16456

Tran, C. H., Vaiana, M., Nakuci, J., Somarowthu, A., Goff, K. M., Goldstein, N., … Goldberg, E. M. (2020). Interneuron Desynchronization Precedes Seizures in a Mouse Model of Dravet Syndrome. The Journal of Neuroscience: The Official Journal of the Society for Neuroscience, 40(13), 2764–2775. https://doi.org/10.1523/JNEUROSCI.2370-19.2020

Traub, R. D., & Miles, R. (1991). Neuronal Networks of the Hippocampus. https://doi.org/10.1017/CBO9780511895401

Ulbricht, W. (2005). Sodium channel inactivation: Molecular determinants and modulation. Physiological Reviews, 85(4), 1271–1301. https://doi.org/10.1152/physrev.00024.2004

Vilin, Y. Y., & Ruben, P. C. (2001). Slow inactivation in voltage-gated sodium channels: Molecular substrates and contributions to channelopathies. Cell Biochemistry and Biophysics, 35(2), 171–190. https://doi.org/10.1385/CBB:35:2:171

Xie, Y., Ng, N. N., Safrina, O. S., Ramos, C. M., Ess, K. C., Schwartz, P. H., … O’Dowd, D. K. (2020). Comparisons of dual isogenic human iPSC pairs identify functional alterations directly caused by an epilepsy associated SCN1A mutation. Neurobiology of Disease, 134, 104627. https://doi.org/10.1016/j.nbd.2019.104627

Yamada, W. M. (1989). Multiple channels and calcium dynamics. Methods in Neuronal Modeling, 97–133.

Yao, Z., Nguyen, T. N., van Velthoven, C. T. J., Goldy, J., Sedeno-Cortes, A. E., Baftizadeh, F., … Zeng, H. (2020). A taxonomy of transcriptomic cell types across the isocortex and hippocampal formation Zizhen. BioRxiv, 10(2), 3.

Yu, F. H., Mantegazza, M., Westenbroek, R. E., Robbins, C. A., Kalume, F., Burton, K. A., … Catterall, W. A. (2006). Reduced sodium current in GABAergic interneurons in a mouse model of severe myoclonic epilepsy in infancy. Nat Neurosci, 9(9), 1142–1149. https://doi.org/10.1038/nn1754

Zuberi, S. M., Brunklaus, A., Birch, R., Reavey, E., Duncan, J., & Forbes, G. H. (2011). Genotype – phenotype associations in SCN1A -related epilepsies. Neurology.

