## Supplemental figures and tables for "Dravet variant *SCN1A^A1783V^* impairs interneuron firing predominantly by altered channel activation"

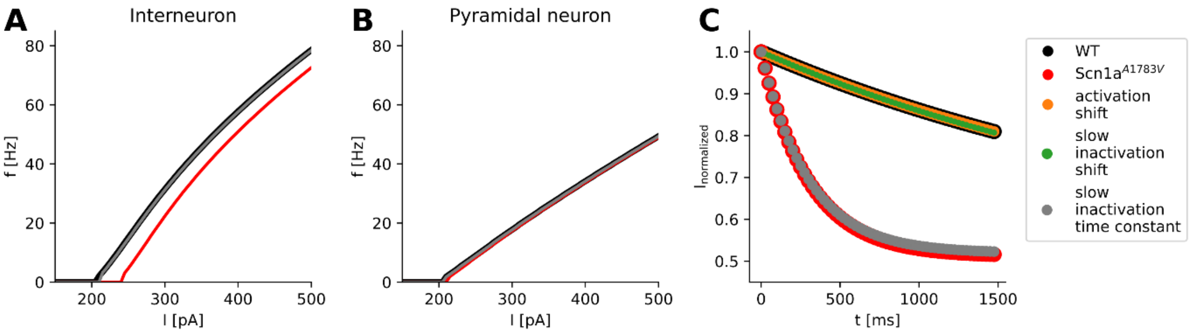


**Supplement figure 1. Simulation study of accelerating the time constant of slow inactivation τ and sodium channel use dependence kinetics.** F-I curves of modelled cortical (A) interneurons and (B) pyramidal cells after acceleration of time constant of slow inactivation *τ* (gray) in comparison to WT (black) and variant *$Scn1a^{A1783V}$* (red) based on findings in tsA cell recordings. (C) Simulated use dependence of sodium current in a cell model solely expressing $Na_V1.1$. at 40 Hz after applying observed changes of activation (orange), slow inactivation (green), time constant *τ* of slow inactivation (grey) and *$SCN1A^{A1783V}$* (red) in comparison to WT (black).

**Supplement table 1: Biophysical properties of *SCN1A* wildtype and mutant channel *$SCN1A^{A1783V}$* recorded in tsA201 cells**

| | **Wildtype** | *n* | **A1783V** | *n* | p |
|---|---|---|---|---|---|
| **Current density [pA/pF]** | -282.2 ± 64.4 | 13 | -284.1 ± 48.57 | 24 | 0.9815 |
| **Steady-state of activation** | | | | | |
| $V_{1/2}$ [mV] | -16.72 ± 2.26 | 13 | -9.17 ± 1.49 | 24 | 0.0066 |
| $k_V$ | -7.794 ± 1.02 | 13 | -7.61 ± 0.31 | 24 | 0.8259 |
| **Fast inactivation** | | | | | |
| $V_{1/2}$ [mV] | -61.31 ± 1.38 | 13 | -61.1 ± 0.80 | 22 | 0.8863 |
| $k$ | 5.54 ± 0.36 | 13 | 5.63 ± 0.37 | 22 | 0.8817 |
| $\tau_{rec}$ at -100 mV [ms] | 6.17 ± 0.57 | 13 | 5.30 ± 0.45 | 14 | 0.24 |
| $\tau_h$ at 0 mV [ms] | 0.34 ± 0.02 | 13 | 0.26 ± 0.02 | 24 | 0.0064 |
| **Slow inactivation** | | | | | |
| $V_{1/2}$ [mV] | -57.93 ± 2.28 | 9 | -68.44 ± 1.23 | 10 | 0.0006 |
| $k$ | 6.07 ± 0.78 | 9 | 4.64 ± 0.31 | 10 | 0.0931 |
| $\tau_{entry}$ [ms] | 3321 ± 220.4 | 12 | 669.1 ± 38.12 | 16 | < 0.0001 |

**Supplement table 2: Electrophysiological properties of cortical fast spiking interneurons recorded in organotypic slice cultures from wildtype and heterozygote floxed Scn1a$^{A1783V}$ mice transduced with AAV8-Syn-Cre-GFP**

| | **Wildtype** | *n* | **A1783V** | *n* | p |
|---|---|---|---|---|---|
| Resting membrane potential [mV] | -69.47 ± 0.72 | 16 | -70.58 ± 1.03 | 16 | 0.3895 |
| Input resistance [MΩ] | 107.8 ± 8.71 | 16 | 73.22 ± 8.87 | 16 | 0.0092 |
| Rheobase [pA] | 209.4 ± 39.52 | 16 | 360.9 ± 40.05 | 16 | 0.012 |
| Maximum firing frequency [Hz] | 88.67 ± 6.00 | 16 | 57.66 ± 9.2 | 16 | 0.0083 |
| AP threshold [mV] | -25.62 ± 3.22 | 13 | -30.3 ± 2.64 | 14 | 0.2686 |
| AP amplitude [mV] | 49.65 ± 3.11 | 13 | 47.86 ± 2.74 | 14 | 0.6684 |
| AP rise time [ms] | 0.75 ± 0.03 | 13 | 0.65 ± 0.04 | 14 | 0.0919 |
| AP half width [ms] | 0.92 ± 0.05 | 13 | 0.78 ± 0.06 | 14 | 0.1168 |

**Supplement table 3: Electrophysiological properties of cortical pyramidal cells recorded in organotypic slice cultures from wildtype and heterozygote floxed Scn1a$^{A1783V}$ mice transduced with AAV8-Syn-Cre-GFP**

| | **Wildtype** | *n* | **A1783V** | *n* | p |
|---|---|---|---|---|---|
| Resting membrane potential [mV] | -73.65 ± 1.398 | 15 | -75.25 ± 1.28 | 25 | 0.424 |
| Input resistance [MΩ] | 263.6 ± 18.3 | 15 | 220.2 ± 11.32 | 25 | 0.040 |
| Rheobase [pA] | 46.67 ± 6.843 | 15 | 50 ± 6.61 | 25 | 0.742 |
| Maximum firing frequency [Hz] | 37.25 ± 3.172 | 15 | 31.7 ± 2.53 | 25 | 0.183 |
| AP threshold [mV] | -44.18 ± 0.907 | 15 | -43.58 ± 1.40 | 25 | 0.762 |
| AP amplitude [mV] | 69.01 ± 3.326 | 15 | 69.03 ± 3.82 | 25 | 0.997 |
| AP rise time [ms] | 1.279 ± 0.055 | 15 | 1.277 ± 0.06 | 25 | 0.987 |
| AP half width [ms] | 1.973 ± 0.147 | 15 | 1.972 ± 0.13 | 25 | 0.996 |
